# Genomic Potential vs. Pathogenic Reality: Functional Virulence Variation in *Pseudomonas aeruginosa*

**DOI:** 10.64898/2026.10.26.761861

**Authors:** R. Dehaene, J. Poppeliers, M. Boon, L. Bens, M. Voet, C. Lood, S. Verwimp, V. Van Noort, G. De Meester, A. Van Daele, A. Crabbé, R. Lavigne

## Abstract

Despite the rapid expansion of bacterial genome sequencing, predicting pathogen virulence from genomic information remains a major challenge. To study this association, we integrate comparative genomics, functional phenotyping, host-pathogen interaction assays, and infection outcomes across a diverse collection of 28 *Pseudomonas aeruginosa* isolates, representing clinically relevant diversity in serotype, type IV pilus group, and flagellar type.

While virulence-associated systems were broadly retained across the strain collection, isolates displayed substantial variation in phenotypic behavior and pathogenicity. Genotypic classifications based on pilus, flagellar, siderophore, or secretion system characteristics were poor predictors of virulence-associated phenotypes. Pathogenicity in a *Galleria mellonella* model varied markedly among strains and was most strongly associated with surface-associated motility, whereas other virulence traits and quorum-sensing genotypes showed limited predictive value. In a three-dimensional lung epithelial cell model, inflammatory responses were negatively correlated with activity of the type III secretion system effector ExoS, suggesting a role in modulation of host immune signaling.

Together, these findings demonstrate that virulence emerges from functional deployment of conserved virulence repertoires, rather than their mere presence. By linking genomic diversity to phenotypic expression, pathogenicity, and host responses within a single strain collection, this study highlights fundamental limitations of gene-content-based virulence prediction and underscores the importance of integrating genomic and functional data to better understand *P. aeruginosa* virulence.

**Importance:** The increasing availability of bacterial genome sequences raises expectations that pathogen virulence can be predicted from gene content. However, despite encoding similar repertoires of virulence-associated genes, many *P. aeruginosa* isolates exhibit markedly different infection outcomes. This study demonstrates that substantial variation in pathogenicity and host interaction can arise from strains with broadly conserved virulence systems. These findings highlight the important contribution of regulatory and functional variation to virulence and show that genomic information alone provides an incomplete picture of pathogenic potential. They further underscore the need to better understand how genomic variation translates into phenotypic traits and host interactions. By integrating genomic analyses with phenotypic characterization and host-response measurements, this work provides the type of dataset needed to connect genomic variation to functional outcomes. Such datasets may ultimately provide the foundation for more reliable prediction of virulence from genome sequences.

## Introduction

Despite extensive genomic characterization, predicting virulence across *Pseudomonas aeruginosa* isolates remains challenging. This limitation reflects the complex and combinatorial nature of the bacterium’s pathogenicity, which arises from the coordinated activity of multiple virulence systems rather than from individual determinants alone (1–3).

Virulence in *P. aeruginosa* depends on the interplay of diverse functional traits, including structural components such as lipopolysaccharide (LPS), motility, secretion systems, and iron acquisition pathways (3–5) (**Error! Reference source not found.**). Flagella-and type IV pili (T4P)-mediated motility supports host colonization and initial surface attachment, and contributes to early biofilm formation, while mature biofilms promote persistence and tolerance to host defenses (6–8). The type II secretion system (T2SS) exports proteases such as elastase (LasB) and protease IV (PrpL) that degrade host tissues and facilitate nutrient acquisition, whereas the type III secretion system (T3SS) injects effector proteins directly into host cells to manipulate cellular processes (9). Among these effectors, ExoS interferes with host signaling pathways and cytoskeletal dynamics, contributing to cellular dysfunction and cytotoxicity (10). In addition, iron acquisition via siderophores such as pyoverdine (PVD) supports bacterial fitness and the regulation of virulence-associated pathways (11). These systems are tightly controlled by regulatory networks, including quorum sensing (QS) (2, 3).

Despite broad retention of these virulence elements across the species, *P. aeruginosa* populations exhibit genetic diversity that arises from allelic variation and differences in accessory genomic elements (12). This diversity leads to distinct LPS serotypes, T4P groups, and flagellar types, some of which are enriched among clinical isolates. LPS serotypes O11, O6 and O1 are the most prevalent in hospital settings, with O1 associated with poor clinical outcomes (13, 14). Furthermore, serotype O12 has been linked to antibiotic resistance (14). Similarly, T4P group I is dominant among clinical isolates, whereas group II is more frequently associated with cystic fibrosis (CF) strains in particular (15, 16). By contrast, flagellar types A1, A2, and B are relatively evenly distributed across clinical populations (14). However, these classifications alone have limited power to predict virulence-associated phenotypes or pathogenicity, suggesting that genomic variability represents only one component of the virulence landscape.

A major challenge in the prediction of virulence is that isolates frequently encode similar sets of virulence-associated genes, while exhibiting different phenotypes *in vitro* and *in vivo* (1, 17, 18). This discrepancy arises from variation in regulatory control, environmental context, and functional gene expression, which are not captured by gene presence alone and can lead to deployment of distinct pathogenic strategies by closely related strains (3, 19, 20).

Although studies of individual virulence determinants have provided important mechanistic insights into *P. aeruginosa* pathogenesis, much of our understanding of virulence regulation and host interaction has been derived from a limited number of laboratory reference strains, particularly PAO1 and PA14 (21). These model strains have been instrumental in defining key virulence pathways, yet they do not fully capture the extensive genetic and phenotypic diversity observed among clinical isolates (21–23). While recent studies have increasingly applied whole-genome sequencing to characterize strain collections and identify virulence-associated genes (12), and others have examined variability in virulence-associated phenotypes across diverse isolates (22, 23), few have systematically integrated genomic diversity, phenotypic behavior, pathogenicity, and host responses within the same strain collection. As a consequence, it remains incompletely understood how genomic diversity translates into virulence-associated phenotypes, pathogenicity, and host responses.

In this study, an integrated analysis of 28 *P. aeruginosa* isolates selected to represent clinical diversity was performed, combining genomic, phenotypic, and host interaction data (**Error! Reference source not found.**). Key virulence traits motility, PVD production, T2SS-dependent protease activity (PrpL and LasB), and T3SS-mediated effector activity (ExoS), were systematically assessed and their contribution to pathogenicity was investigated in a *Galleria mellonella* infection model. For a subset of eight strains, host responses were investigated using a three-dimensional (3D) A549 lung epithelial cell model, to determine how bacterial virulence profiles translate into immunogenic outcomes. By linking genomic potential, phenotypic behavior, and host interaction, this study investigates how conserved *P. aeruginosa* repertoires give rise to divergent pathogenic outcomes and provides insight into the complex factors that shape virulence expression across diverse strains.

## Results

### *In silico* virulence analysis across a *P. aeruginosa* collection representing clinically relevant diversity

To enable assessment of correlations between virulence and genomic features, the genomic diversity of key virulence systems within a collection of isolates was examined. From the large collection of *P. aeruginosa* isolates previously characterized by Pirnay et al. (18), 28 strains were selected to capture clinically relevant diversity (Table 1). Specifically, selection was based on serotype (O1, O6, O11, O12), T4P group (I and II), and flagellar type (A1, A2, B), which together represent the most prevalent and clinically relevant characteristics reported for *P. aeruginosa* (13–16). For each of the four serotypes, clinical non-CF strains were selected to capture variation in T4P and flagellar types. Where available, CF, environmental, and animal isolates were added to maximize representation of the selected genotypes, which occur across a range of ecological and host-associated settings. PAO1, although a serotype O2 isolate, was added to the collection as a reference strain.

**Table 1.** Genomic characteristics of the Pseudomonas aeruginosa strain collection. Strains are shown with their source of isolation, ST, LPS serotype, T4P group, flagellar e, T2SS effector *toxA* status, T3SS effectors profile, PVD siderovar and QS system integrity. All strains were obtained from the collection of *P. aeruginosa* isolates previously racterized by Pirnay et al. (18).

| Strain | Source | ST | Serotype | T4P group | Flagellar type | T2SS *toxA* | T3SS effector | Siderovar | Quorum Sensing systems |
| --- | --- | --- | --- | --- | --- | --- | --- | --- | --- |
| LMG25175 | Medical - Wound | 313 | O1 | 1 | A2 | *toxA* | *exoTUY* | I | **No *lasR*** |
| LMG24967 | Medical - Pleural fluid | 3893 | O1 | 2 | A2 | *toxA* | ***exoSTUY*** | I | **No *lasRI*,**<br>**C-term. trunc. *rhlR*, No *pqsR*** |
| LMG25036 | Medical - CF-patient | 787 | O1/O6* | 1 | A1 | *toxA* | *exoSTY* | II | Intact |
| LMG25010 | Medical - CF-patient | 845 | O1 | 2 | A1 | *toxA* | *exoSTY* | II | Intact |
| LMG25025 | Medical - CF-patient | 17 | O1 | **(2)** | A1 | *toxA* | *exoSTY* | II | Intact |
| LMG24972 | Environmental - Tobacco plant | 1920 | O1 | 2 | B | *toxA* | *exoSTY* | I | **C-term. trunc. *pqsB*** |
| LMG25181 | Animal - Dog eye | 1400 | O1 | 1 | A2 | *toxA* | *exoTUY* | I | **C-term. trunc. *lasR*** |
| LMG25053 | Medical - Throat | 233 | O6 | 1 | A1 | *toxA* | *exoSTY* | II | **No *lasR*** |
| LMG25078 | Medical (non-CF) | 7003 | O6 | 2 | B | *toxA* | *exoSTY* | III | **C-term. trunc. *pqsE*** |
| LMG25069 | Medical - CF-patient | 395 | O6 | 1 | A1 | *toxA* | *exoSTY* | I | Intact |
| LMG25058 | Medical - CF-patient | 2035 | O6 | 2 | A2 | *toxA* | *exoSTY* | II | Intact |
| LMG25044 | Environmental - River water | 179 | O6 | 1 | A1 | *toxA* | *exoSTY* | II | Intact |
| LMG24998 | Environmental - Lake water | 876 | O6 | 2 | A1 | *toxA* | *exoSTY* | II | Intact |
| LMG25105 | Environmental - River water | NA | O6 | 2 | A1 | *toxA* | *exoSTY* | II | **No *lasRI*** |
| LMG25201 | Animal - Dog | 1716 | O6 | 2 | B | **NA** | **NA** | II | Intact |
| LMG25124 | Medical - Burn | 671 | O11/O7* | 1 | B | *toxA* | *exoTUY* | II | Intact |
| LMG24891 | Medical - Surgical bandage | **4646** | O11 | 2 | A1 | **NA** | ***exoST*** | III | Intact |
| LMG25194 | Medical - Wound | **4646** | O11 | 2 | A1 | **NA** | ***exoST*** | III | Intact |
| LMG25156 | Medical - Sputum | **446** | O11 | 2 | A2 | *toxA* | *exoTUY* | I | Intact |
| LMG24887 | Medical - Sputum | 2045 | O11 | 2 | B | *toxA* | *exoSTY* | III | Intact |
| LMG25148 | Hospital environment | 308 | O11 | 2 | A2 | *toxA* | *exoTUY* | II | Intact |
| LMG25164 | Environmental - Ocean water | 316 | O11 | 2 | A2 | *toxA* | *exoTUY* | I | **No *lasR*** |
| LMG25173 | Environmental - River water | **446** | O11 | 2 | A2 | *toxA* | *exoTUY* | I | **C-term. trunc. *lasR*** |
| LMG24889 | Animal - Dog | NA | O11 | 2 | A2 | *toxA* | *exoSTY* | III | **No *lasRI*** |
| LMG24912 | Medical (non-CF) | **111** | O12 | 1 | B | *toxA* | *exoSTY* | III | Intact |
| LMG24926 | Medical - Sputum | **111** | O12 | 1 | B | *toxA* | *exoSTY* | III | Intact |
| LMG25200** | Medical - Wound | 1195 | O12 | 2 | A2 | **NA** | **NA** | II | Intact |
| PAO1 | Medical - Wound | 549 | O2 | 2 | B | *toxA* | *exoSTY* | I | Intact |

\*Strains where serotyping by slide agglutination (before /) and serotyping using the pasty tool (after /) rendered different results.
\*\*Strain LMG25200 or PA7 was reclassified as species *Pseudomonas paraaeruginosa* in 2022 (24).

Notably, strain LMG25200 or PA7 was reclassified in 2022 as *Pseudomonas paraeruginosa*, a closely related member of the *P. aeruginosa* species complex (24). It was nevertheless retained in the collection due to its historical inclusion among *P. aeruginosa* isolates and its relevance in comparative analyses and clinical contexts. To verify the classification of the remaining strains, a rapid whole-genome similarity assessment was performed using FastANI (Galaxy v1.3) (25). All strains show an ANI of ≥97.5% relative to PAO1, whereas ANI values do not exceed 94% when compared to LMG25200, supporting the latter’s distinct taxonomic status within the collection (Table S1).

After establishment of the strain collection, long-read whole-genome sequencing and annotation were performed for all 28 strains to improve assembly contiguity, resulting in less fragmented genomes compared to the Illumina-only assemblies. For subsequent genomic analyses of the strain collection, Nanopore-derived genome assemblies were used (Table S2). MLST analysis assigned sequence types (STs) to 26 out of the 28 strains. Among these, 23 unique STs were identified, with three pairs of strains sharing the same ST (ST111, ST446, and ST4646) (Table 1).

To obtain a global overview of the virulence repertoire present within the strain collection, all 28 genomes were screened for virulence-associated genes (26). Virulence determinants are broadly conserved across the collection, differing only in the absence of T3SS-associated genes in two strains. Next, four virulence factors of interest were selected for further investigation within the strain collection: (1) motility, (2) PVD production, (3) T2SS, and (4) T3SS. These systems were selected because they capture complementary aspects of *P. aeruginosa* virulence, including host colonization (motility), nutrient acquisition (PVD), extracellular tissue damage (T2SS-associated proteases), and direct host-cell manipulation (T3SS effectors), while also permitting quantification using *in vitro* bio-assays. Prior to phenotypic characterization, the genomic basis of these factors was assessed across all strains. This included evaluation of completeness of the virulence-associated systems, as well as their classification.

### Genomic assessment reveals overall completeness of virulence system

All four systems assessed are present and largely complete in terms of their structural and regulatory genes across the strain collection, except for the T3SS, which is confirmed to be absent in two strains (LMG25200 and LMG25201). This finding is expected for *P. paraeruginosa* strain LMG25200, as this species generally lacks the T3SS and its associated effectors. In contrast, strain LMG25201 is classified as *P. aeruginosa,* but does share some characteristics with the *P. paraeruginosa* strain. Notably, these two strains are the only ones in the collection to encode the exolysin ExlA, a pore-forming virulence factor typically associated with *P. (par)aeruginosa* strains that lack the classical T3SS and that employ ExlA-mediated cytotoxicity as an alternative strategy for host-cell damage (27, 28).

Despite overall completeness, some strains lack individual components. Strains LMG25025 and LMG25124 lack essential components of the T4P and PVD clusters, respectively. Given the high completeness of the genome assemblies, these absences are unlikely to reflect assembly artefacts. More specifically, in strain LMG25025, *pilM* was not detected by TBLASTN, despite its identification by TXSScan, suggesting sequence divergence or a false-negative. PilM is a component of the PilMNOP pilus alignment subcomplex, which links the outer membrane secretin with the inner membrane motor subcomplexes. This highly conserved complex is responsible for positioning the pilus machinery and contributes to both extension and retraction (29). Disruption of any of the subcomplex components results in loss of surface-exposed pili (30). This is expected to result in the loss of twitching motility and impairment of several T4P-dependent functions, including adhesion to host cells, microcolony formation and surface sensing (6, 31). Notably, strain LMG25025 originates from a CF infection. Chronic CF isolates commonly exhibit reduced motility and adaptation towards a biofilm-associated lifestyle, suggesting that loss of specific virulence-associated functions may reflect niche adaptation and persistence within the CF lung rather than generalized attenuation (32). For strain LMG25124, assessment of the antiSMASH-identified PVD cluster reveals the absence of core genes *pvdI* and *pvdJ*, indicating loss of PVD production. Additionally, in strain LMG24889, the PVD cluster is split into two separate regions due to integration of a prophage. However, as all core biosynthetic genes remain intact, this insertion does not necessarily imply disruption of PVD production.

Alongside the structural and regulatory elements, effector gene repertoires of the T2SS and T3SS were also examined. For the T2SS, most effector genes (*lasA, lasB, prpL, plcH, plcN*) are retained across the collection, whereas *toxA*, encoding Exotoxin A, is absent in four strains (LMG24891, LMG25194, LMG25200, LMG25201) (Table 1). Previous research has shown that *toxA* is not universally present in clinical *P. aeruginosa* strains, reflecting variability in virulence gene repertoires (33). Notably, three of the four *toxA*-negative strains are of a medical, non-CF source, further indicating that Exotoxin A is not essential for pathogenicity in all contexts.

For the T3SS, effector profiling reveals the expected distribution of *exoS* and *exoU*, which are typically mutually exclusive and associated with different infection characteristics (34–36). Interestingly, one strain (LMG24967) encoded both *exoS* and *exoU*, with the two genes annotated at separate genomic loci and distinguishable based on sequence identity (threshold 75%), supporting their independent presence rather than a detection artefact. While this co-occurrence is atypical, it has been documented previously in distinct *P. aeruginosa* isolates, including several internationally disseminated high-risk clones. The *exoS+/exoU+* genotype has been associated with increased virulence and is likely facilitated by horizontal gene transfer, as *exoU* is commonly located on mobile genomic islands (37–39). In contrast, IslandViewer4 (40) does not predict a genomic island at the *exoU* locus of LMG24967. The same isolate was previously classified as *exoS*+/*exoU*-based on PCR analysis (18). However, inspection of the *exoU* locus confirms that both published *exoU* primer binding sites are present, and no genomic explanation for the earlier negative PCR result could be identified. An additional discrepancy with previous PCR-based typing can be observed for strain LMG25194, which is found to carry *exoS* despite previously being reported as negative (18). Analysis of the primer annealing regions shows that the reverse primer binding site, located downstream of *exoS* in the adjacent intergenic region, is absent, providing a likely explanation for the previous false-negative result and highlighting a limitation of PCR-based genotyping approaches that rely on conserved primer-binding regions.

Beyond variation in the *exoS/exoU* repertoire, two strains (LMG24891 & LMG25194) lack *exoY*, further illustrating that, despite broad conservation of the structural T3SS machinery, effector repertoires can differ considerably between strains (Table 1) (35).

### Virulence systems segregate into distinct genomic groups

Next, the virulence systems were classified into distinct genotypic groups where possible. For motility, this classification was already established, with strains categorized into T4P groups based on the major pilin *pilA* and associated genes (group I (n = 9 strains); group II (n= 19 strains)), and into flagellar types based on the major flagellin gene *fliC* (type A1 (n = 10 strains); type A2 (n= 10 strains); type B (n= 8 strains)).

Based on variation in their PVD biosynthetic clusters, strains were classified into three phylogenetically distinct types, termed siderovars. More specifically, classification was based on the *fpvA* gene encoding the ferripyoverdine receptor, as variation within the biosynthetic cluster is reflected in corresponding changes in the receptor sequence (41, 42). Consistent with Pirnay et al. (18), this results in a distribution of nine type I, twelve type II, and twelve type III strains, reflecting balanced representation of all three types within the collection (Table 1).

In contrast, analysis of the T2SS and T3SS gene clusters revealed little sequence or compositional variation among strains. Classification based on either gene cluster composition or hallmark genes (T2SS: *xcpQ, xcpT, xcpQ*; T3SS: *pscF, pcrV, pscN*) did not separate strains, reflecting the high conservation of the core secretion machinery (9). Consequently, for both secretion systems, strain differentiation is only possible at the level of the previously identified differences in effector profiles.

In conclusion, the four selected virulence systems are broadly retained across the strain collection, while variation is primarily observed in system-specific genotypic groups and effector profiles. Strains are classified based on T4P groups, flagellar types, T2SS and T3SS effector profiles, and PVD-based siderovars. In a next step, phenotypic virulence traits were assessed, allowing functional differences between strains to be interpreted in light of the genomic features identified above.

### Functional characterization of *P. aeruginosa* virulence traits

Given the broad conservation of core virulence systems, functional characterization of the *P. aeruginosa* strain collection was performed using *in vitro* bio-assays targeted to each of the four selected virulence factors (**Error! Reference source not found.**). This was complemented by evaluation of all 28 strains in the *in vivo G. mellonella* infection model to assess overall virulence in a host context.

### Targeted bio-assays show no association with previously defined genomic groups

All targeted bio-assays revealed extensive variability across the strain collection. Substantial differences can be observed in motility, PVD production, and secretion system activity, with no consistent patterns across strains (Table 2).

**Table 2.**
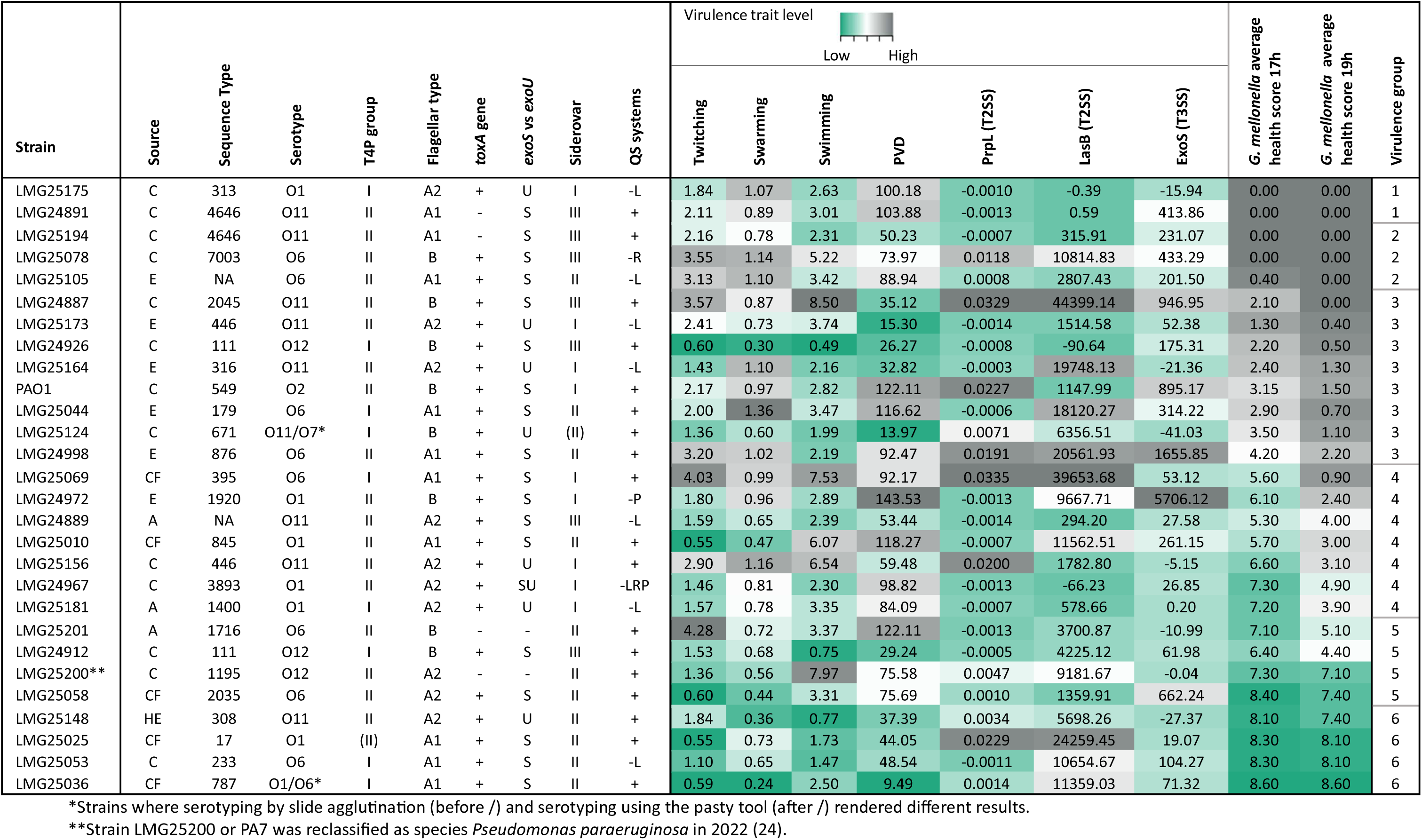
Integrated heatmap of virulence-associated phenotypes across the *P. aeruginosa* strain collection. Heatmap showing quantitative results for all *in vitro* virulence assays, including motility (swimming, swarming, twitching; diameters [cm]), PVD production (relative fluorescence [a.u.]), T2SS activity (PrpL (2.5*(A405/time)) [min^-1^])) and LasB (F415/time) [min^-1^]), and T3SS activity (ExoS (F410/time) [min^-1^]), together with *G. mellonella* health scores at 17 h and 19 h post-infection. Color reflects relative activity levels per virulence trait (green = least virulent; grey = most virulent). Annotation tracks indicate genomic features for each strain, including source (C = Clinical non-CF, CF = Clinical CF, E = Environmental, A = Animal, HE = Hospital Environment), ST, serotype, T4P group, flagellar type, T2SS and T3SS effector profiles, PVD siderovar and QS system completeness (+ = complete; - = potentially impaired; L= Las, R = Rhl, P = PQS). Strains are arranged according to increasing virulence. Virulence groups (1 = most virulent, 6 = least virulent) were defined based on pairwise Kaplan-Meier survival comparisons (log-rank test with Benjamini-Hochberg correction).

For motility, variability is evident across swimming, swarming, and twitching assays, with strains displaying distinct and often assay-specific behaviors, not associated to T4P types or flagellar groups (Figure S1). Strain LMG25025, for which no *pilM* gene was annotated, indeed exhibits the lowest twitching diameter among all strains tested, consistent with the predicted disruption of the PilMNOP alignment subcomplex and the consequent loss of functional T4P.

Similarly, PVD production shows a wide range of outputs when measuring PVD-associated fluorescence, with no clear association with the assigned siderovars (Figure S2). As expected, strain LMG25124 lacking *pvdIJ,* exhibits markedly low PVD-associated fluorescence. For both the motility and PVD production traits, strains sharing the same genotypic classification frequently exhibit markedly different phenotypes.

A high degree of heterogeneity can also be observed for both secretion systems. T2SS activity, assessed through PrpL and LasB production, varies considerably between strains, and shows discordant behavior in several isolates, with high production of one substrate but not the other. Although both enzymes are secreted via the T2SS, their expression is regulated by distinct pathways, with LasB primarily controlled by QS, and PrpL mainly regulated by iron availability via PvdS. This allows differential expression under varying environmental conditions (43–45). Despite these differences in transcriptional regulation, the activities of both proteases are interconnected, as LasB contributes to the post-secretional activity of PrpL by cleaving its inhibitory propeptide (46, 47). However, it should be noted that PrpL and LasB were quantified under different assay conditions optimized for expression of each protease (see Materials & Methods), which may have further contributed to the observed variation in their production among the strains.

T3SS activity, measured by ExoS secretion, displays a skewed distribution, with several strains exhibiting little to no activity and only a limited subset showing high expression. As expected, strains lacking an annotated *exoS* gene do not secrete any ExoS. However, the absence of secretion is not restricted to this group, as several *exoS*-positive strains also exhibit little to no detectable ExoS production. This shows that presence of effector genes is not sufficient to predict their production. T3SS activity is tightly regulated through the ExsA-ExsD-ExsC-ExsE regulatory cascade, which integrates environmental signals to control expression of T3SS genes (48, 49). Consequently, variation in regulatory control, differences in T3SS induction, or defects affecting secretion system assembly may result in little to no detectable ExoS production despite the presence of an intact *exoS* gene.

Overall, the *in vitro* assays revealed substantial heterogeneity in virulence factor production across the strain collection, highlighting the strong regulatory influences and condition-dependent expression. Under the tested conditions, previously defined genomic classifications showed little predictive value for virulence-associated phenotypes (data not shown), whereas absence of core genes reliably indicated loss of the corresponding virulence factor. Notably, this heterogeneity extends even to strains sharing the same ST. Although the ST4646 pair exhibits highly similar phenotypes, the two other pairs show significant differences in motility (ST11 & ST446) and PrpL production (ST446), illustrating that strains considered clonally related may nevertheless exhibit markedly different phenotypic behavior. As these assays evaluate individual factors, they do not capture their combined contribution during infection. Therefore, all strains were further evaluated in the *G. mellonella* infection model to assess their overall virulence.

**Divergent pathogenicity in the *Galleria mellonella* model is not captured by overall *in vitro*** virulence profiles

Consistent with the *in vitro* findings, substantial variability in virulence can be observed in the *G. mellonella* infection model. Kaplan–Meier survival analysis reveals significant differences in pathogenicity between strains, with some causing rapid host mortality and others exhibiting limited virulence (Figure S3 & Table S3). These differences enable classification of the bacterial strains into six virulence classes (1 = most virulent; 6 = not virulent) (Table 2). While highly virulent strains are already distinguishable at 15 h, the greatest divergence in larval health occurs at 17-19 h. Health scores within this interval were therefore used as a quantitative measure of virulence for integration with *in vitro* data (Table 2).

No obvious association can be observed between pathogenicity in the *G. mellonella* model and previously characterized *in vitro* virulence characteristics (Table 2). This lack of concordance is consistent with different *P. aeruginosa* strains employing distinct combinations of virulence-associated traits to establish infection. This was further illustrated by strains sharing the same ST. While the ST4646 pair exhibited largely similar phenotypes and were both among the most pathogenic strains in the collection, the ST111 and ST446 pairs differed not only in motility and PrpL production but also in pathogenicity in the *G. mellonella* model. Within the ST111 pair, the more motile strain, LMG24912, as expected displayed higher pathogenicity than LMG24926. In contrast, within the ST446 pair, LMG25173 exhibited higher virulence despite lower motility and PrpL production than LMG25156. Possible associations between the different virulence assays were further investigated using correlation analysis.

### Motility-associated phenotypes correlate with virulence in the *Galleria mellonella* model

Pairwise Spearman correlations were calculated across all assays (Figure 2A & Table S4). A strong correlation can be observed between larval health scores at 17 h and 19 h (*ρ* = 0.97, *p* < 0.0001), confirming the robustness of the selected time window (Figure 2B).

**Figure 1.**
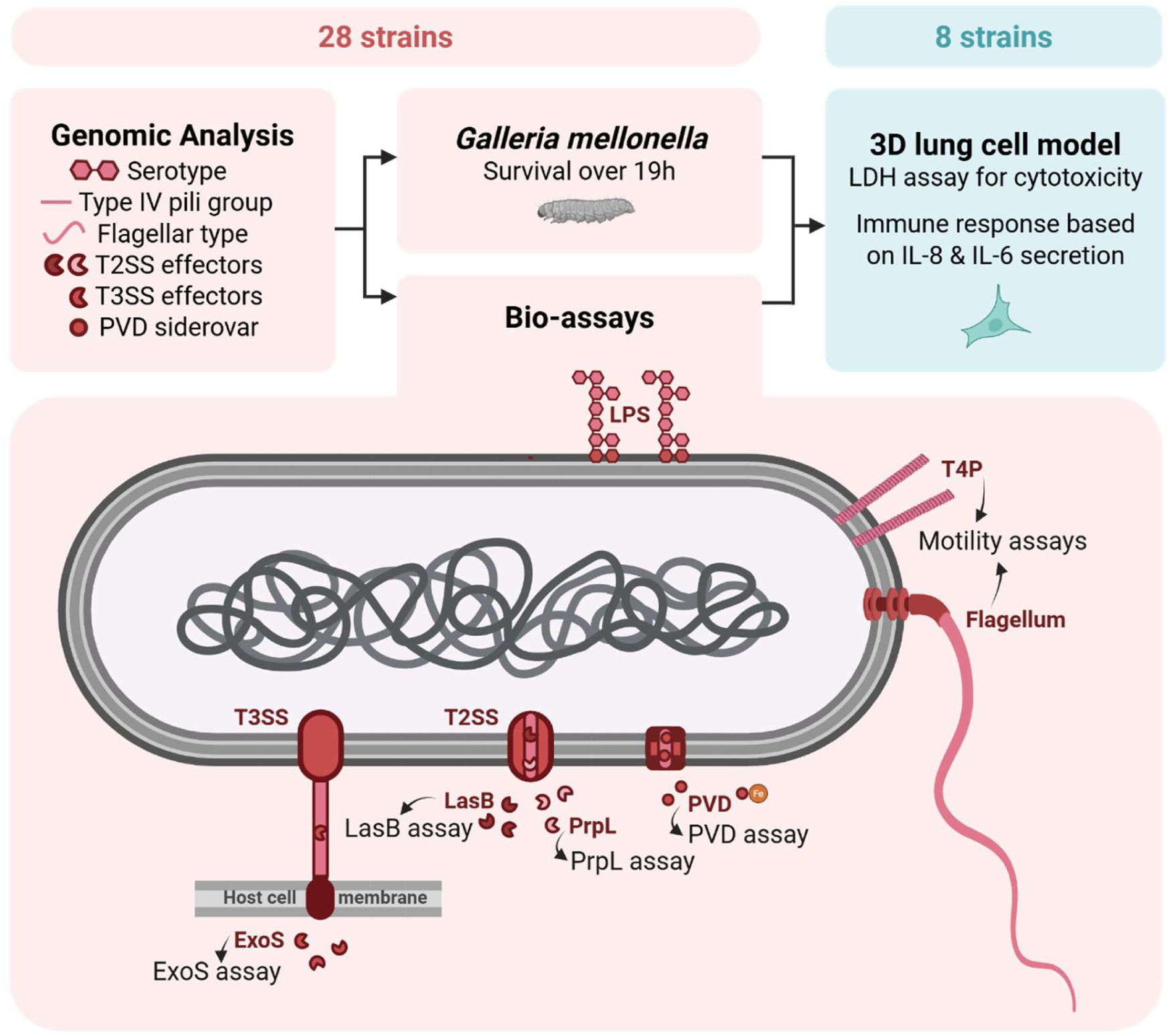
Integrated workflow for virulence profiling of the *P. aeruginosa* strain collection. 28 *P. aeruginosa* strains were subjected to genomic characterization, virulence-associated bio-assays, and infection studies in *G. mellonella*. Genomic features analyzed included serotype, T4P group, flagellar type, T2SS and T3SS effector repertoires, and PVD siderovar (top left). Based on their combined bio-assay and *G. mellonella* virulence profiles (middle), eight virulent strains were selected for further evaluation in a 3D lung epithelial cell model (top right). The lower panel illustrates the virulence-associated systems examined and the corresponding functional assays used in this study.

**Figure 2.**
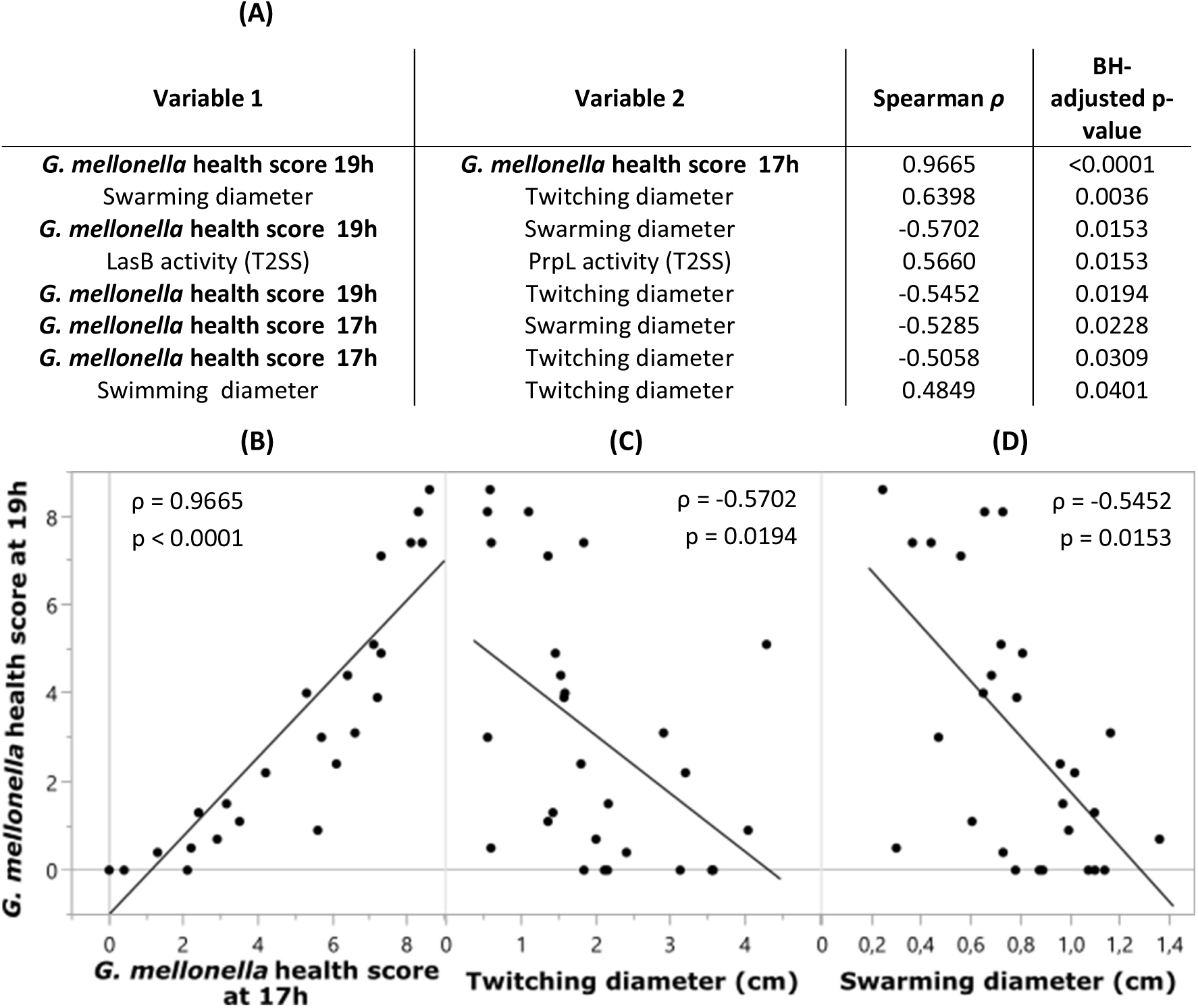
Associations between phenotypic traits and virulence in the *G. mellonella* model. (A) Significant pairwise Spearman rank correlations between phenotypic assays (BH-adjusted *p*-value < 0.05), ordered by increasing BH-adjusted *p*-value. (B) Scatterplot comparing larval health scores at 17 h and 19 h. (C) Scatterplot of twitching motility (diameter) vs larval health scores at 19 h. (D) Scatterplot of swarming motility (diameter) vs larval health scores at 19 h. Each point represents an individual strain. Spearman correlation coefficients *ρ* and corresponding BH-adjusted *p*-values are indicated in the scatterplots. Lines represent linear trends for visualization only.

Correlation analysis further identified expected associations between phenotypic traits. Despite the previously described discordant production patterns in several strains, a significant positive correlation can be observed between the T2SS-associated proteases LasB and PrpL (*ρ* = 0.57, *p* = 0.0153), consistent with their shared secretion pathway and post-secretion interactions (46, 47). Motility assays are strongly associated, with twitching correlating with both swarming (*ρ* = 0.64, *p* = 0.0036) and swimming motility (*ρ* = 0.48, *p* = 0.0401).

Among the *in vitro* virulence phenotypes, motility-associated traits show the strongest associations with pathogenicity in the *G. mellonella* model. Both swarming and twitching motility are significantly negatively correlated with larval health scores at 17 h (swarming: *ρ* =-0.57, *p =* 0.0228; twitching: *ρ* =-0.55, *p* = 0.0309) and 19 h (swarming: *ρ* =-0.53, *p* = 0.0153; twitching: *ρ* =-0.51, *p* = 0.0194), indicating that increased motility is associated with higher virulence (Figure 2C-D). This is interesting because it highlights motility as a determinant of pathogenic potential in the *G. mellonella* model, suggesting that isolates with enhanced surface-associated movement are better able to establish infection and cause host damage.

While motility shows association with virulence in the *G. mellonella* model, this factor still only partially explains the observed variability. The identified patterns of trait co-occurrence further suggest that virulence is shaped by coordinated expression of multiple phenotypes. Given that many of these traits are under QS control, the contribution of QS system variation to phenotypic and virulence differences was examined.

### Genomic QS variation shows limited association with phenotypic variation and virulence

Genomic variation in QS systems was identified across the strain collection (Table 1), including the absence or C-terminal truncation of LasR(I) in several strains. Furthermore, strains LMG24972 and LMG25078 contain truncations in the PQS operon, specifically in PqsB (C-term, 33 aa) and PqsE (C-term, 60 aa), respectively. Whereas disruption of PqsB is expected to impair PQS biosynthesis, truncation of PqsE likely affects Rhl-dependent regulation, since PqsE modulates Rhl-controlled gene expression (50, 51). Notably, strain LMG24967 displays extensive QS deficiency, lacking LasR and PqsR and harboring a severely truncated RhlR (C-term, 38aa), indicative of a near-complete breakdown of hierarchical QS regulation.

Despite the observed impairments, correlations between QS genotypes and phenotypic outputs are limited. No consistent associations are observed for motility, PVD production or ExoS activity, although that one strain with putative PQS impairment (LMG24972) shows elevated PVD and ExoS, consistent with known effects of PQS on iron regulation and T3SS cytotoxin secretion (52–54) (Figure 3A). Similarly, variability in the *G. mellonella* model does not correspond to QS genotypes, with strains harboring Las deficiencies displaying divergent virulence phenotypes (Figure 3B).

**Figure 3.**
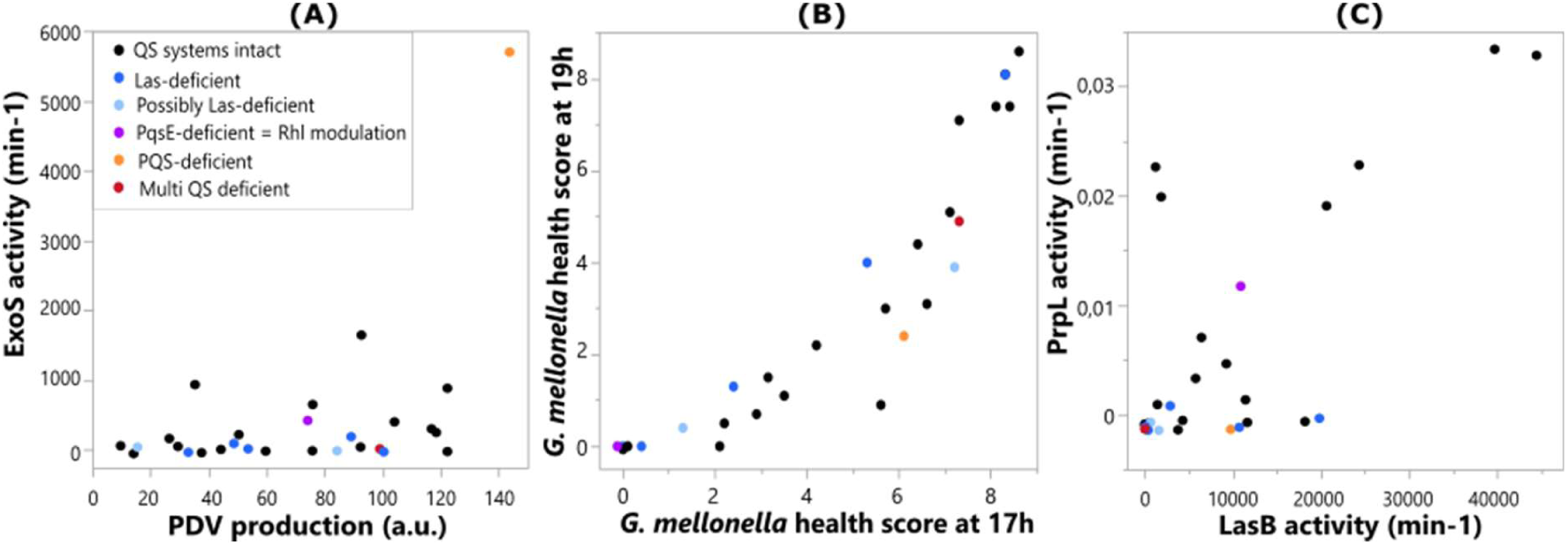
Associations between phenotypic traits and QS genotypes across the strain collection. Scatterplots show (A) PVD production vs ExoS activity, (B) *G. mellonella* larval health scores at 17 h vs 19 h post-infection, and (C) PrpL vs LasB production. Each point represents an individual strain and is colored according to QS genotype classification.

A potential association between Las impairment and reduced PrpL secretion can be observed, although this is not consistently mirrored by reduced LasB production (Figure 3C). This unexpected pattern, given their respective regulatory controls, may indicate indirect effects of QS disruption on PrpL expression or secretion, but could equally result from variation in iron-responsive regulatory pathways independent of LasR status (45). Furthermore, LasB contributes to the post-secretional activation of PrpL by cleaving its inhibitory propeptide. However, elevated LasB levels cannot compensate for low PrpL expression (46, 47). Consequently, reduced PrpL activity in LasR-impaired isolates should be interpreted as the result of interactions between QS, iron-responsive regulation, and strain-specific adaptive changes. Notably, LasR impairment does not consistently translate to loss of LasB activity, suggesting that some LasR-deficient strains partially compensate for the loss of canonical Las signaling. Previous studies have demonstrated that clinical *P. aeruginosa* isolates can undergo QS rewiring, allowing maintenance of QS-regulated virulence traits through alternative regulatory circuits involving RhlR and PqsE (50, 55–57). More broadly, differential evolutionary adaptions to QS defects are likely to contribute to the pronounced phenotypical heterogeneity observed across strains (58).

Overall, these findings indicate that while QS systems influence specific aspects of virulence, their genomic background alone does not provide a comprehensive explanation for the observed phenotypic variability, nor is it predictive of pathogenic potential.

### Strain-dependent epithelial inflammatory responses in a 3D alveolar lung model

To investigate whether virulence potential translates into host immune cell activation, a subset of eight strains was selected for evaluation in a physiologically relevant 3D alveolar epithelial model based on A549 cells. Compared with conventional monolayer cultures, this model more closely reflects the structural organization and functional properties of the alveolar epithelium, supporting epithelial polarization and barrier formation, while maintaining biologically relevant cytokine responses following inflammatory stimulation (59–61). Strain selection was directed toward high virulence isolates, defined by strong expression of multiple virulence-associated traits and high pathogenicity in the *G. mellonella* model. Reference strain PAO1 was included as a benchmark and also exhibited a high virulence profile.

To confirm that differences in downstream cytokine production are not confounded by variation in host cell viability, cytotoxicity was assessed at 4 h post-infection. No significant differences were observed between infected conditions and the uninfected control, indicating that none of the strains induced cytotoxicity within this timeframe and changes in cytokine production reflected genuine differences in host cell activation (Figure S4).

### ExoS activity is inversely associated with epithelial cytokine responses

Quantification of cytokine production reveals pronounced strain-dependent variation in cytokine responses (Table 3). Both interleukin (IL)-8 and interleukin (IL)-6 production vary substantially across strains, with a strong concordance between the two markers. This coordinated behavior is consistent with activation of shared inflammatory pathways downstream of *P. aeruginosa-*host interactions (62–64). Contrary to our initial hypothesis, strains exhibiting the lowest pathogenicity in the larval infection model appear to elicit the strongest cytokine responses in the epithelial system. In line with this observation, cytokine production shows a moderate positive Spearman correlation with *G. mellonella* health scores, particularly at 17 h post infection (IL-6: *ρ* = 0.57; IL-8: *ρ* = 0.52) (Figure 4A&C), indicating that virulence in the *G. mellonella* model does not necessarily translate into stronger epithelial immune activation.

**Figure 4.**
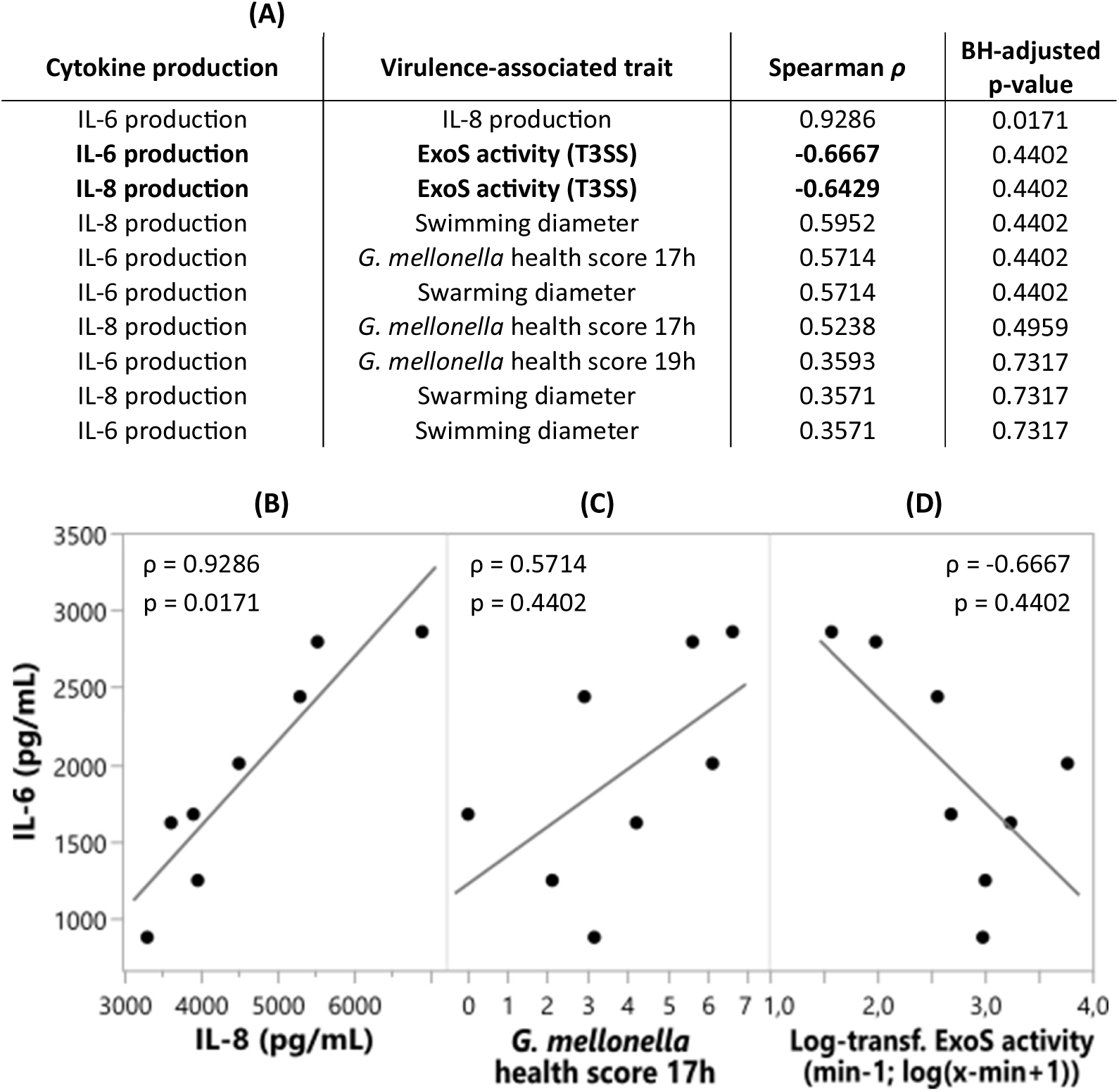
Associations between epithelial cytokine responses and virulence-associated traits. (A) Pairwise Spearman rank correlations between cytokine responses and virulence-associated traits, ordered by increasing BH-adjusted *p*-value. Only correlations with a BH-adjusted *p*-value < 0.8 are shown. (B) Scatterplot comparing IL-8 and IL-6 production after 4 h exposure of the 3D alveolar epithelial model. (C) Scatterplot of IL-6 production vs larval health scores at 17 h post-infection in the *G. mellonella* model. (D) Scatterplot of IL-6 production vs log-transformed ExoS activity. Each point represents an individual strain. Spearman correlation coefficients (*ρ*) and corresponding BH-adjusted *p*-values are indicated in the scatterplots. Lines represent linear trends for visualization purposes only.

**Table 3.**
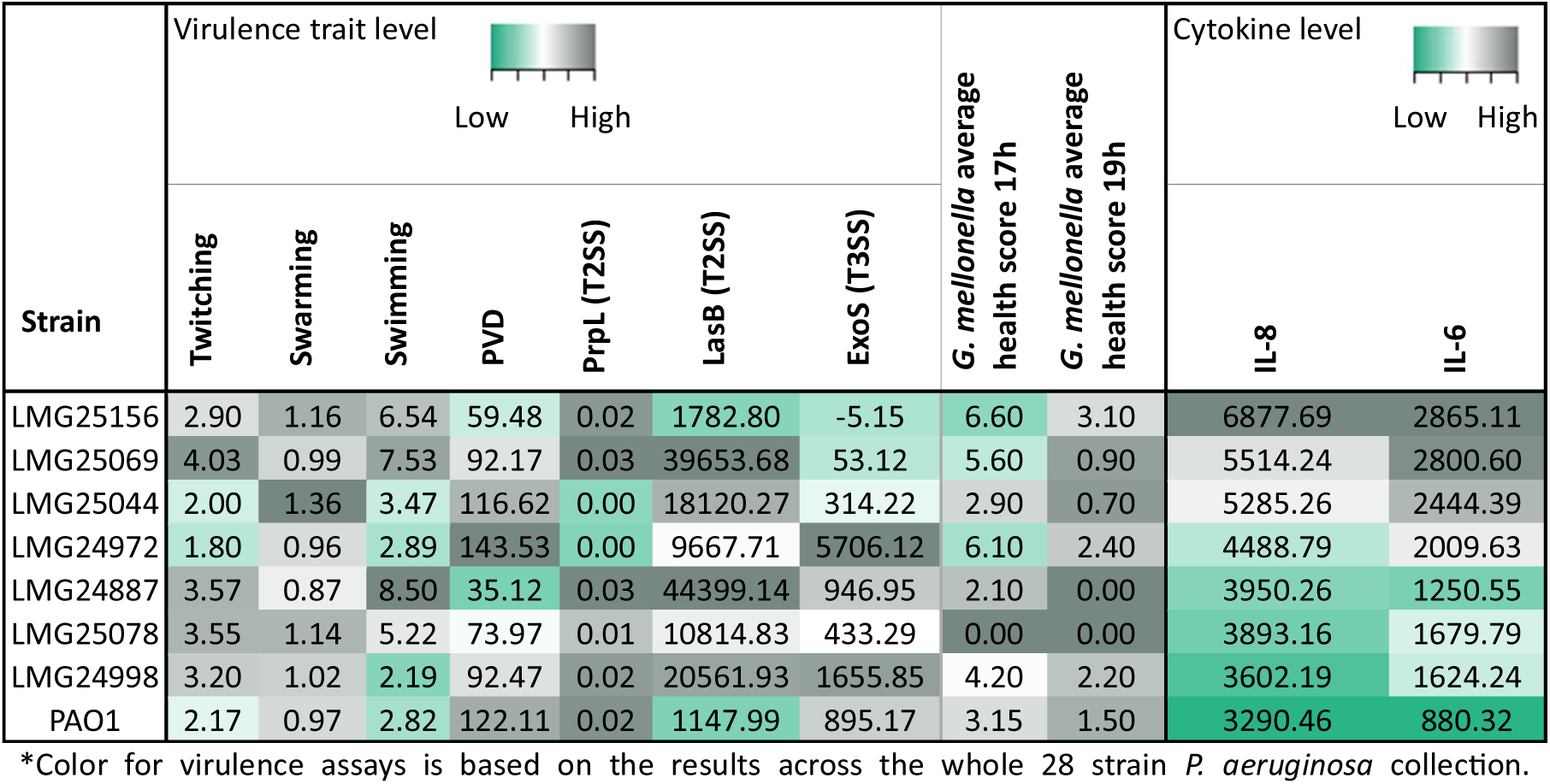
Integrated heatmap of virulence-associated phenotypes and epithelial cytokine responses for 8 selected *P. aeruginosa* strains tested in the 3D lung cell model. Heatmap showing quantitative results for all *in vitro* virulence assays, including motility (swimming, swarming, twitching; diameters [cm]), PVD production (relative fluorescence [a.u.]), T2SS activity (PrpL (2.5*(A405/time)) [min⁻¹] and LasB (F415/time) [min⁻¹]), and T3SS activity (ExoS (F410/time) [min⁻¹]), together with larval health scores at 17h and 19 h post-infection in the *G. mellonella* model and epithelial cytokine production (IL-6 and IL-8 [pg/mL]) after 4 h exposure in the 3D alveolar lung model. Strains are ordered based on increasing cytokine production, and color reflects relative activity levels per assay (green = low; grey = high).

A potential explanation for this observation may not lie with overall pathogenicity, but with differences in T3SS effector ExoS. To investigate this possibility, Spearman rank correlations were calculated between all virulence assays and cytokine production (Figure 4A & Table S5). IL-8 and IL-6 levels show a strong positive correlation (*ρ* = 0.93, *p =* 0.0171) (Figure 4B). While sample size (n=8) limits statistical power and definitive conclusions, ExoS activity displays the strongest and most consistent association with cytokine production among the virulence-associated traits examined. Both IL-8 and IL-6 show moderate negative correlations with ExoS (IL-8: *ρ* =-0.64, *p* = 0.4402; IL-6: *ρ* =-0.67, *p* = 0.0442), hinting towards lower cytokine production in strains with higher ExoS activity (Figure 4D). Although the underlying mechanism remains unclear, this observation is consistent with the established ability of ExoS to modulate host inflammation (65). While these findings are based on a limited number of strains, they identify ExoS as a potential determinant of variation in epithelial immune responses, suggesting that pathogenicity does not only depend on the expression of virulence factors but also on the ability to modulate host inflammation.

Other virulence-associated traits, including motility phenotypes, also exhibit moderate positive correlations with cytokine production (*ρ* ≈ 0.57–0.60), but these associations are less consistent across assays and cytokines, and do not reveal a clear pattern.

## Discussion

This study provides a comprehensive characterization of virulence-associated traits across a collection of 28 *P. aeruginosa* isolates selected to represent clinically relevant diversity, by integrating genomic analyses, functional phenotyping, infection model outcomes, and host-cell responses. Despite the broad genomic retention of core virulence-associated systems across the collection, substantial heterogeneity could be observed at the phenotypic level and strain classifications based on allelic variability were poor predictors of virulence factor production. This disconnect was evident even among strains sharing the same ST.

Among the phenotypes examined, swarming and twitching motility show significant associations with pathogenicity in the *G. mellonella* model. Isolates exhibiting enhanced surface-associated motility generally induce greater larval mortality, suggesting that these motility phenotypes may serve as indicators of pathogenic potential in this model. This observation is consistent with previous studies showing that T4P and flagella contribute not only to bacterial movement, but also to host colonization, surface sensing, and interactions with host immune defenses (6, 7, 66). Furthermore, swarming motility has been associated with coordinated expression of multiple virulence determinants, including secretion systems, proteases, and iron acquisition pathways (67). While most virulence traits did not correlate significantly with swarming in the present study, a moderate positive trend was observed between swarming motility and PVD production (ρ = 0.4291, *p* = 0.0908). Together with the association between swarming and pathogenicity observed here, this supports the idea that motility may reflect a broader virulence-associated physiological state rather than acting as an isolated determinant of pathogenicity. Importantly, associations between motility and pathogenicity have also been demonstrated in mammalian infection models, where functional T4P and twitching motility contribute to epithelial interactions, host colonization, cytotoxicity, and dissemination, whereas swarming-associated behavior enhances bacterial spread and fitness during infection (68–70). Extending these observations, our results show that natural variation in swarming and twitching phenotypes across a diverse strain collection is associated with differences in pathogenicity. This trend was further reflected within the ST111 pair, where the more motile strain also exhibited higher pathogenicity. However, motility only partly explains the observed variation in virulence. Notably, the ST446 pair deviated from this trend, as LMG25173 displayed higher pathogenicity despite reduced motility and PrpL production compared to LMG25156, indicating that additional determinants contribute to pathogenicity. Together, these observations illustrate that different isolates employ distinct combinations of virulence mechanisms to establish infection and cause host damage.

Comparison of pathogenicity and host inflammatory responses reveals an additional level of complexity. Strains displaying lower pathogenicity in *G. mellonella* induce higher IL-6 and IL-8 production in a 3D A549 lung epithelial model, an observation associated with reduced ExoS activity. Although based on a limited subset of isolates, these findings suggest that ExoS may contribute to modulation of epithelial inflammatory responses. Interestingly, this negative correlation observed between ExoS activity and epithelial IL-6/IL-8 production contrasts with previous observations in peripheral blood mononuclear cells (PBMCs), where purified ExoS induced a strong pro-inflammatory cytokine response, including elevated IL-6 and IL-8 levels (71). This discrepancy may reflect differences in host cell type, the use of extracellular purified ExoS versus bacterially delivered ExoS, or context-dependent effects of ExoS on immune signaling. Furthermore, more recent studies have demonstrated that ExoS can suppress host defense pathways, including neutrophil ROS production and inflammasome-associated responses (72–74), supporting a broader role for ExoS in immune suppression as opposed to the pro-inflammatory effect of the T3SS-structural components (65). Together, these findings suggest that the impact of ExoS on host immunity may be dependent on host cell type and infection context and warrant further investigation.

From a translational perspective, these findings not only have implications for pathogenicity prediction, but also for anti-virulence strategies. The identification of swarming and twitching as virulence-associated phenotypes linked to pathogenicity suggests that pathways governing surface-associated motility warrant further investigation as potential anti-virulence targets. Given the established roles of T4P and flagella in host colonization, surface sensing, tissue interaction, and dissemination during mammalian infection (6, 7, 66, 68–70), interference with motility-associated processes could affect multiple stages of the infection process. Furthermore, previous reports linking swarming to coordinated expression of multiple virulence determinants (67) raise the possibility that targeting motility-associated pathways may influence broader virulence-associated physiological states rather than bacterial movement alone. Consequently, strategies aimed at disrupting motility-related processes may represent an approach to reduce pathogenicity without directly affecting bacterial viability, thereby potentially limiting selective pressure for resistance development. In addition, a better understanding of how ExoS and the T3SS shape host inflammatory responses may provide new insights into host-pathogen interactions and identify targets for therapies aimed at restoring protective immune responses during infection. Further work will be required to determine whether epithelial cytokine modulation is directly mediated by ExoS or reflects broader alterations in T3SS activity.

Collectively, this study supports the view that *P. aeruginosa* virulence represents a complex and combinatorial phenotype that emerges from the coordinated activity of multiple virulence systems, regulatory networks and host-pathogen processes, rather than from individual determinants in isolation (1–3). Strains sharing similar virulence gene repertoires or genomic classifications frequently displayed markedly different phenotypes and pathogenic outcomes, highlighting the limitations of predicting virulence from gene content alone and underscoring the importance of regulatory mechanisms and environmental influences in shaping virulence expression. Notably, substantial phenotypic divergence was observed even among strains sharing the same ST, emphasizing that close genetic relatedness does not necessarily translate into similar virulence-associated behavior or pathogenic potential. These observations point to two important implications for future research. First, integrating transcriptomic, proteomic, and functional datasets will be required to identify the regulatory mechanisms driving the disconnect between virulence gene content, phenotypic behavior and pathogenicity. Ultimately, resolving these genotype-phenotype relationships may enable more accurate prediction of virulence from genome sequences. Second, incorporating diverse strain collections rather than relying solely on a small number of laboratory reference strains for virulence studies, infection models and therapeutic development pipelines, will improve the translational relevance of experimental findings and provide a more realistic representation of the pathogenic strategies employed by clinical *P. aeruginosa* populations.

## Materials & Methods

### Bacterial cultures & growth conditions

Bacterial cultures (Table 1) were streaked from glycerol stocks onto 1.5% lysogeny broth (LB) agar plates. Unless otherwise stated, overnight cultures were grown in LB at 37 °C with shaking.

### Strain characterization & genomic analysis

Out of the large *Pseudomonas aeruginosa* collection previously characterized by Pirnay et al. (18), 28 strains were selected based on LPS serotypes, T4P group, and flagellar type. For 322 out of 328 strains, Illumina whole-genome sequencing data was available (75).

### LPS serotyping

Previously, LPS serotypes were determined by slide agglutination. However, only 215 out of the 328 strains could be serotyped in this manner (18). To complement the data, *in silico* serotyping was performed using pasty (v1.0.3), which assigns O-antigen serogroups by detecting and comparing O-specific antigen biosynthesis loci in genome assemblies against a reference database using sequence similarity searches (76, 77).

### T4P grouping

T4P grouping is based on phylogenetically distinct groups of pilin gene *pilA* alleles, each linked to a characteristic set of accessory genes (15). A genomic context-based approach was used to identify candidate T4P loci and assign groups based on the detected gene combinations.

First, T4P-associated genes were identified using a custom nucleotide BLAST-based approach. A curated query dataset was compiled containing representative sequences of key T4P-associated genes, including multiple *pilA* allelic variants, *pilB*, and accessory genes (*tfpOa*, *tfpOb*, *tfpW*, *tfpX*, *tfpY*, *tfpZ*) (Table S6). Genome assemblies in fasta format were screened against this query set using BLASTN (BLAST+ suite, v2.15.0) (78). Searches were performed using an *e*-value threshold of 0.05 and a minimum query coverage of 30% per high-scoring pair (HSP).

tRNA genes were predicted using tRNAscan-SE (v2.0.12) (79, 80) with bacterial search mode enabled. Only tRNA-Thr loci were retained. Candidate T4P loci were defined as genomic regions in which a *pilB* gene and a tRNA-Thr locus co-occurred on the same contig within a maximum distance of 20 kb.

Within each candidate region, the presence or absence of additional T4P-associated genes was assessed based on BLAST hits fully contained within the defined interval. Based on the combination of detected genes, isolates were classified into the five T4P groups: Group I encoding *pilA* + *tfpOa/b*, Group II encoding *pilA* only, with no additional T4P-associated genes detected in the region between the tRNA-Thr locus and *pilB,* Group III encoding *pilA* + *tfpY,* Group IV encoding *pilA* + *tfpW* and *tfpX* and finally, Group V encoding *pilA* + *tfpZ* (15). Only BLAST hits fully contained within the candidate region were considered for group assignment.

### Flagellar typing

Flagellar typing was performed using *fliC* gene sequences identified in all available Prokaryotic Genome Annotation Pipeline (version 2024-07-18.build7555) (81) annotated genomes. Differentiation between major flagellar types A and B was based on the length of the *fliC* gene, where isolates with *fliC* gene lengths of 1164 bp or 1185 bp were classified as type A, whereas those with longer gene lengths of 1458 bp or 1467 bp were classified as type B (82). Further subtyping within type A was determined by examining specific deletions in the derived amino acid sequence. Specifically, subtype A2 is characterized by two deletions of 3 and 4 amino acids at positions 254 and 279, respectively, relative to the A1 reference sequence from strain PAK (83).

### Nanopore-based whole-genome sequencing

The DNeasy UltraClean Microbial Kit (Qiagen; Hilden, Germany) was used for genomic DNA extraction following the manufacturer’s protocol. Whole-genome sequencing was performed on extracted genomes using the PromethION device (Oxford Nanopore Technologies; Oxford, UK). Further analysis was performed with an automated snakemake pipeline. Basecalling was performed using Dorado v1.1.1 (Nanopore). Next, raw sequencing data was cleaned by trimming off barcodes with porechop_abi v0.5.1 (84) and selecting reads with a quality score above Q10 using chopper v0.11.0 (85). The resulting data was used for genome assembly with Flye v2.9.6 (86). Finally, the assembled genome was annotated using Bakta v1.11.4 (87).

### Multilocus sequence typing

MLST was performed on Nanopore-based genome assemblies using the MLST tool (Galaxy version 2.22.0) (88). For strains for which automatic ST assignment was unsuccessful, STs were manually annotated by comparing the allelic profile against the PubMLST database (89).

### Genomic assessment of virulence system completeness

Genome assemblies were screened for virulence-associated genes using ABRicate (v1.0.1) (26) against the VFDB database (90). Hits were retained at minimum nucleotide identity and reference sequence coverage thresholds of 80%. Following this screening, completeness of virulence-associated systems was assessed using a combination of automated detection and manual sequence-based verification.

For the T4P, flagellar system, T2SS and T3SS, an initial screening was performed using TXSScan (v1.1.4) (91). For each predicted system, TXSScan reports a completeness score (1 = complete gene set), together with the detected protein components (mandatory and accessory) and their corresponding genomic coordinates. Notably, TXSScan does not assess completeness of the full flagellar assembly pathway, as its flagellum model is limited to components sufficient to distinguish it from T3SS.

To complement this analysis, tBLASTn (BLAST+ suite, v2.17.0+) (78) searches were conducted using manually curated query sets for each individual system (Table S7). Searches were performed with an *e*-value threshold of 1 × 10⁻³ and a minimum query coverage per high-scoring pair (HSP) of 70% The query sets comprised structural and regulatory genes with known or predicted roles in the functioning of the respective virulence systems. For T2SS and T3SS, effector proteins were additionally included. The gene sets were used to verify the presence of the complete gene repertoire in each genome.

The tBLASTn approach was also extended to assess completeness of the *P. aeruginosa* QS systems Las, Rhl and PQS, for which a separate manually curated query set was compiled (Table S7).

To assess PVD pathway completeness, secondary metabolite biosynthetic gene clusters were identified using antiSMASH (v8.0.4) (92). Completeness of the PVD system was subsequently assessed by evaluating the presence of the core biosynthetic genes (*pvdL, pvdIJD, pvdO, pvdP, pvdA)* within each identified PVD cluster.

### Virulence assays

#### Semi-quantitative pyoverdine detection assay

To evaluate PVD production, 2 μL of overnight culture was spotted onto solid CAA agar plates (Per liter: 5 g acid-hydrolyzed casein, 1.18 g K_2_HPO_4_, 1 mL mgSO_4_ [1M] and 15 g bacterial agar). For each strain, three biological replicates were plated on CAA without FeCl_3_ (stimulation of PVD production), and three replicates were plated on CAA supplemented with 150 μM FeCl_3_ (negative control). Plates were incubated at 37 °C for 6h.

PVD fluorescence was visualized using a G:BOX imaging system equipped with an UltraBright-Blue LED transilluminator (Syngene) and images were captured (Iris = 1.6, Zoom = 18.3, Focus = 100). Image analysis was performed using Fiji (93). Circular regions of interest (ROIs, diameter = 50 pixels) were defined around each colony, and the mean grey value was measured. Background correction was applied by subtracting the average mean grey value of five background ROIs from each measurement.

#### Motility assays

Motility was assessed by measuring swimming, swarming and twitching diameters.

For the swimming and swarming assays, overnight cultures were used to inoculate the center of agar plates using a sterile toothpick. Swimming and swarming were evaluated on 0.3% and 0.5% LB agar plates, respectively. For swimming motility, plates were incubated at 37 °C for 15 h 30 min, after which the diameter of the spreading zone was measured. For swarming motility, plates were incubated at 30 °C for 24h, before measuring the spreading diameters. For both assays, three biological replicates were performed per strain.

To evaluate twitching motility, an adapted version of the macroscopic twitching assay as described by Turnbull and Whitchurch was performed (94). Briefly, overnight cultures were stab-inoculated to the bottom of 1% LB agar plates using a sterile toothpick. Plates were incubated at 37 °C for 48 h. Subsequently, the agar was removed and the plate bottoms were stained using TM developer solution (Per liter: 400 mL water, 100 mL glacial acetic acid, 500 mL methanol; stored at 4 °C) to visualise the interstitial colony at the agar-plastic interface. The solution was left to dry for 15 min and then decanted. The diameter of the stained twitching zone was measured. Five biological replicates were performed per strain.

### PrpL detection assay

To determine the activity of the proteinase IV enzyme (PrpL) secreted into the supernatant by the T2SS, the assay described by Massai et al. (95) was adapted. In short, overnight cultures were grown in CAA medium (Per liter: 5 g acid-hydrolyzed casein, 1.18 g K_2_HPO_4_ and 1 mL mgSO_4_ (1M)) supplied with 150 μM FeCl_3_, at 37 °C. The following day, cultures were diluted 1:500 in fresh CAA medium without FeCl_3_, to induce PrpL secretion. The diluted cultures were incubated for 16h at 37 °C with shaking.

After incubation, OD_600nm_ was measured and culture supernatant was collected. Supernatants were diluted 2.5-fold, and 20 μL was mixed with 60 μL phosphate buffer (Na_2_HPO_4_ (0.05M) and NaH_2_PO_4_ (0.05M); pH 7) and 20 μL Chromozym PL (2 mg/mL; Merck). Protease activity was monitored by measuring absorbance at 405 nm every two min for 30 min at 25 °C using a CLARIOstar Plus microplate reader. A minimum of three biological replicates was performed per strain.

### LasB detection assay

To quantify the activity of the elastase enzyme (LasB) secreted into the supernatant by the T2SS, a fluorimetric assay of peptidase activity was performed as described by Kessler & Safrin (96–98). Briefly, overnight cultures were prepared and culture supernatant was collected and diluted 1:100 in assay buffer (Tris–HCl (0.05 M), CaCl_2_ (2.5 mM); pH 7.2).

For the reaction, 10 μL of diluted supernatant was mixed with 90 μL of Abz-Ala-Gly-Leu-Nba (AAGLAN) substrate working solution (110 μM) and 10 μL assay buffer in a black flat-bottom 96-well plate. Elastase activity was monitored by measuring fluorescence (excitation at 330 nm, emission at 415 nm) every two min for 30 min at 25 °C using a CLARIOstar Plus microplate reader. Three biological replicates were performed per strain.

### ExoS detection assay

To assess the activity of the T3SS toxin ExoS in culture supernatant, the protocol by Pinto et al. (99) was adapted. In short, overnight cultures were 100-fold diluted in LBi (for 0.5 L: 450 mL LB, 50 mL EGTA (50 mM) and 10 mL MgCl_2_ (1M)) and incubated at 37 °C with shaking for four hours. After incubation, OD_600nm_ of the cultures was measured and culture supernatant was collected.

The assays were performed in a black, flat-bottom 96-well plate. For the reaction, 20 μL of supernatant was mixed with 2 μL 14-3-3 (500 mM), 2 μLνH-Ras (25 μM) and 71 μL HEPES buffer (HEPES salts (20 mM), NaCl (50 mM), MgCl_2_ (4 mM) and TCEP (0.5 mM); pH 7.5; 4 °C). The reaction was initiated by adding 5 μL NAD^+^ (500 μM). ExoS activity was monitored by measuring fluorescence (excitation at 320 nm, emission at 410 nm) every 2 min for 44 min at 25 °C using a CLARIOstar Plus microplate reader. Three biological replicates were performed per strain.

### Galleria mellonella virulence assay

*In vivo* virulence was assessed using the *G. mellonella* infection model. Ten larvae (300-550 mg) were selected per bacterial strain. Overnight cultures were diluted in PBS (NaCl (137 mM), KCl (2.7 mM), Na_2_HPO_4_ (8.2 mM) and 1.8 mM KH_2_PO_4_; pH 7.4) to +- 1000 CFU/mL. A volume of 10 μL of the diluted bacterial culture or PBS (negative control) was injected in the hindmost right proleg of each larva using a fine needle (Micro-Fine^TM^ Plus U-100 insulin syringes, BD, Le Pont-de-Claix, FR). Following injection, the larvae were kept in a 12-well culture plate (CELLSTAR® 12-well cell culture plates, Greiner Bio-One, Kremsmünster, Austria; cat. no. 665180) and placed in the dark at 37 °C. Larval survival and health status were monitored at 15 h, 17 h and 19 h post-injection.

Larval health was quantified using a scoring matrix (Table 4) based on activity, melanization and survival (100). A maximum score of 9 represents optimal health, whereas dead larvae were assigned a health score of 0.

**Table 4.** Scoring matrix for assessment of larval health (adapted from (100))

| Category | Description | Score |
| --- | --- | --- |
| Activity | No activity | 0 |
|  | Minimal activity on stimulation | 1 |
|  | Active when stimulated | 2 |
|  | Active without stimulation | 3 |
| Melanization | Complete melanization (black) | 0 |
|  | Dark spots on brown wax worm | 1 |
| | ≥ 3 spots on beige wax worm | 2 |
| | $<$ 3 spots on beige wax worm | 3 |
|  | No melanization | 4 |
| Survival | Dead | 0 |
|  | Alive | 2 |
| **Total** | **Sum of all categories** | **/9** |

To compare survival between groups, Kaplan-Meier survival analysis was performed in R (v4.3.3) using the survival and survminer packages within the RStudio environment (Posit Software, PBC, Boston, MA, USA). Survival curves were compared using the log-rank test. Pairwise comparisons between strains were performed using log-rank tests with Benjamini–Hochberg correction for multiple testing. Based on these comparisons, strains were categorized into six virulence classes.

### Host response assays in a 3D alveolar lung model

#### Three-dimensional lung epithelial cell culture model

The human adenocarcinomic alveolar epithelial cell line A549 (ATCC, CCL185) was cultured as an organotypic three-dimensional (3D) cell culture model using the rotating wall vessel (RWV) bioreactor, as previously described (101–103). A549 cells were grown as monolayers in T75 flasks in serum-free GTSF-2 medium (Hyclone™, Logan, UT, USA) supplemented with 1.5 g/L sodium bicarbonate (Sigma-Aldrich), 10% heat-inactivated fetal bovine serum (FBS) (Thermo Fisher Scientific) and2.5 mg/L insulin transferrin sodium selenite (Sigma-Aldrich).. Upon reaching confluence, cells were washed with Hank’s Balanced Salt Solution (HBSS) (Life Technologies, Carlsbad, CA, USA) and detached using 0.25% trypsin-EDTA solution (Thermo Fisher Scientific). Cell numbers and viability were assessed using 0.4% Trypan Blue (Sigma-Aldrich) and a Bürker chamber. A suspension containing 2 × 10^6^ cells in supplemented GTSF-2 medium was then combined with 0.25 g dextran beads coated with porcine-skin collagen (Cytodex®-3 microcarrier beads, Cytiva, Marlborough, MA, USA). The mixture was transferred into a RWV (slow turning lateral vessel (STLV) and high-aspect rotating vessel (HARV) (Synthecon, Houston, TX, USA)), and cultured for 11 to 14 days to allow formation of differentiated 3D lung cell aggregates. All cultures were incubated at 37 °C, 5% CO_2_.

#### *In vitro* cell exposure assay

For each strain, a 5 mL overnight culture was pelleted and the supernatant was discarded. The bacterial pellet was then resuspended in 5 mL serum-free GTSF-2 medium and the OD_600nm_ was measured. The suspension was then diluted in serum-free GTSF-2 medium to a final concentration of 5 x 10^7^ CFU/mL.

3D cell aggregates were transferred to a flat-bottom 96-well plate at a density of 2.5 x 10^5^ cells/well in serum-free GTSF-2 medium. The aggregates (200 μL) were exposed to 50 μL of the bacterial suspension, resulting in a final multiplicity of infection (MOI) (bacteria:A549 alveolar cells) of 10. The 96-well plate was incubated for 4 hours (5% CO_2_, 37 °C). For each bacterial strain and the negative control (GTSF-2 medium), five biological replicates were performed, with each biological replicate analyzed in duplicate (technical replicates).

After the four hour incubation period, 150 μL of supernatant from each infected well was collected and transferred to a new 96-well flat-bottom plate. To prevent cytokine degradation by bacterial proteases, Halt^TM^ Protease Inhibitor Cocktail (100x) and 0.5 M EDTA Solution (Thermo Fisher, Ref. 87786) were each added at a 1:100 dilution (1.5 μL per 150 μL supernatant). The supernatants were stored at-20 °C until further analysis by ELISA.

#### Cytotoxicity assay

To assess cytotoxicity, the protocol developed by Van den Bossche et al. was followed (104). This protocol solves the issue of lactate dehydrogenase (LDH) degradation by *P. aeruginosa* proteases. In short, after removing the cell culture supernatant, the A549 cell aggregates were washed 2x with HBSS. The viable cells were lysed using 1% Triton X-100 (Sigma-Aldrich). Finally, the Lactate Dehydrogenase Activity Assay Kit (Sigma-Aldrich, MAK066) was used to quantify LDH activity per the manufacturer’s instructions.

#### Inflammatory marker quantification

IL-8 and IL-6 release was measured in the cell culture supernatant using the Human ELISA MAX^TM^ *Standard Set* (IL-8 Cat. No. 431501; IL-6 Cat. No. 430501) by Biolegend®, according to the manufacturer’s instructions. Optimal dilutions of the samples were prepared in GTSF-2 medium without FBS.

#### Spearman rank correlations & scatter plots

Pairwise associations between assays were visualized using scatter plots and evaluated using Spearman rank correlation analysis. Scatter plots, correlation coefficients (ρ), and corresponding two-sided *p*-values were generated in JMP® Student Edition 19 (JMP Statistical Discovery LLC, Cary, NC). To account for multiple testing, *p*-values within each correlation analysis were manually adjusted using Benjamini–Hochberg correction. Spearman correlations (*ρ*) with an adjusted *p*-value < 0.05 were considered statistically significant.

## Acknowledgements

The authors thank Prof. Jean-Paul Pirnay for providing access to the *Pseudomonas aeruginosa* strain collection used in this study.

This work was supported by the European Research Council (ERC) under the European Union’s Horizon 2020 Research and Innovation Programme [819800 to R.L.]. M.B. was supported by a grant from the Special Research Fund [iBOF/21/092]. R.L. and V.N. are funded through the KU Leuven project C1 ‘ACES’ (C16/20/001). C.L. was supported by a junior postdoctoral fellowship from the Research Foundation – Flanders (FWO) [12D8623N].

## Supplemental Material

*See doc ‘Dehaene-et-al_mBIO_Supplemental-Material*.

## Data Availability

Raw DNA sequencing files are deposited under NCBI BioProject ID PRJNA1540791. Any additional data is accessible from the authors upon request.

